# Neural entrainment determines the words we hear

**DOI:** 10.1101/175000

**Authors:** Anne Kösem, Hans Rutger Bosker, Atsuko Takashima, Antje Meyer, Ole Jensen, Peter Hagoort

## Abstract

Low-frequency neural entrainment to rhythmic input has been hypothesized as a canonical mechanism that shapes sensory perception in time. Neural entrainment is deemed particularly relevant for speech analysis, as it would contribute to the extraction of discrete linguistic elements from continuous acoustic signals. Yet, its causal influence in speech perception has been difficult to establish. Here, we provide evidence that oscillations build temporal predictions about the duration of speech tokens that directly influence perception. Using magnetoencephalography (MEG), we studied neural dynamics during listening to sentences that changed in speech rate. We observed neural entrainment to preceding speech rhythms persisting for several cycles after the change in rate. The sustained entrainment was associated with changes in the perceived duration of the last word’s vowel, resulting in the perception of words with radically different meanings. These findings support oscillatory models of speech processing, suggesting that neural oscillations actively shape speech perception.

## INTRODUCTION

Brain oscillations are known to entrain to rhythmic sensory signals. Neural entrainment is observed for various stimulation ranges and sensory modalities, yet it is still unclear whether the observed oscillatory activity in electrophysiological recordings truly reflects the recruitment of endogenous neural oscillations, and whether these oscillations causally influence sensory processing and perception [1]. Neural entrainment that relies on the recruitment of endogenous oscillations should be dynamic and self-sustained, meaning that it should adapt to the dynamics of current sensory rhythms and should persist for several cycles after stimulation. Crucially, the sustained neural entrainment would be functionally relevant for sensory processing as it would provide a temporal predictive mechanism [2,3]: neural entrainment would reflect the internalization of past sensory rhythms to optimize sensory processing by predicting the timing of future sensory events. So far evidence for sustained entrainment is scarce, and has only been reported in occipital cortices for visual alpha oscillations, and in temporal cortices after auditory entrainment in monkey recordings [4,5]. A crucial open question is whether sustained entrainment occurs during the presentation of complex ecological signals such as speech, and, if so, how it would impact perception [6,7].

Neural entrainment could provide important temporal information for speech processing, given that the acoustic signal presents periodicities of the same temporal granularity as relevant linguistic units, e.g. syllables [6,7]. Specifically, low-frequency neural entrainment has been proposed to contribute to parsing, and to defining the duration of discrete speech information extracted from the continuous auditory input [8–10]. Being recruited at the earliest stages of speech analysis, entrained oscillations should ultimately influence the perception of the spoken utterances. As for other entrainment schemes, their causal efficacy in speech processing remains debated [11–14]. Because neural oscillations match the dynamics of speech during entrainment, it is unclear whether oscillatory activity observed in electrophysiological recordings during speech processing reflects the involvement of neural oscillators for speech analysis, or, alternatively, is the consequence of non-oscillatory based mechanisms that modulate the evoked response to the rhythmic speech signal [13]. For instance, stronger neural entrainment has repeatedly been observed for more intelligible speech signals [15–18], but these observations could either originate from the stronger recruitment of oscillatory mechanisms, or from the enhanced evoked response to the speech acoustic features.

To demonstrate the causal role of neural entrainment in speech perception, the oscillatory activity has to be disentangled from the driving stimulus’s dynamics. Neural oscillatory models suggest that this dissociation is possible when speech temporal characteristics are suddenly changing. Sustained entrainment to the preceding speech dynamics should be observed after a change in speech rate, meaning that the observed neural entrainment to speech is dependent on contextual rhythmic information. If neural oscillations causally influence speech processing, different neural oscillatory dynamics should lead to different percepts for the same speech material. This predicts that entrainment to past speech rhythms should influence subsequent perception. In line with this proposal, contextual speech rate has been shown to affect the detection of subsequent words [19], word segmentation boundaries [20], and perceived constituent durations [21–23]. We propose that these effects could originate from the presence of sustained neural oscillatory activity that defines the parsing window of linguistic segments from continuous speech [8,13,21]. The frequency of sustained entrainment should then affect the onset, offset and size of the discretized items, so that a change in frequency leads to distinct percepts of the extracted linguistic units.

We tested this hypothesis in an MEG study in which native Dutch participants listened to Dutch sentences with varying speech rates. The beginning of the sentence (carrier window) was either presented at a fast or a slow speech rate (Fig. 1A). Specifically, during the carrier window, the speech envelopes in the slow and fast rate conditions had a strong rhythmic component at 3 Hz and 5.5 Hz respectively (Fig. 1B). The last three words (target window) were consistently presented at an intermediate pace (Fig. 1C). Participants were asked to report their perception of the last word of the sentence (target word), which contained a vowel ambiguous between a short /α/ and a long /a:/, and could be perceived as two distinct Dutch words (e.g., *tak* /tαk/ “branch” or *taak* /ta:k/ “task”). We investigated whether sustained neural entrainment to speech could be visible after a speech rate change (during the target window), and if the sustained entrainment causally affected the perception of the target word.

**Figure 1:**
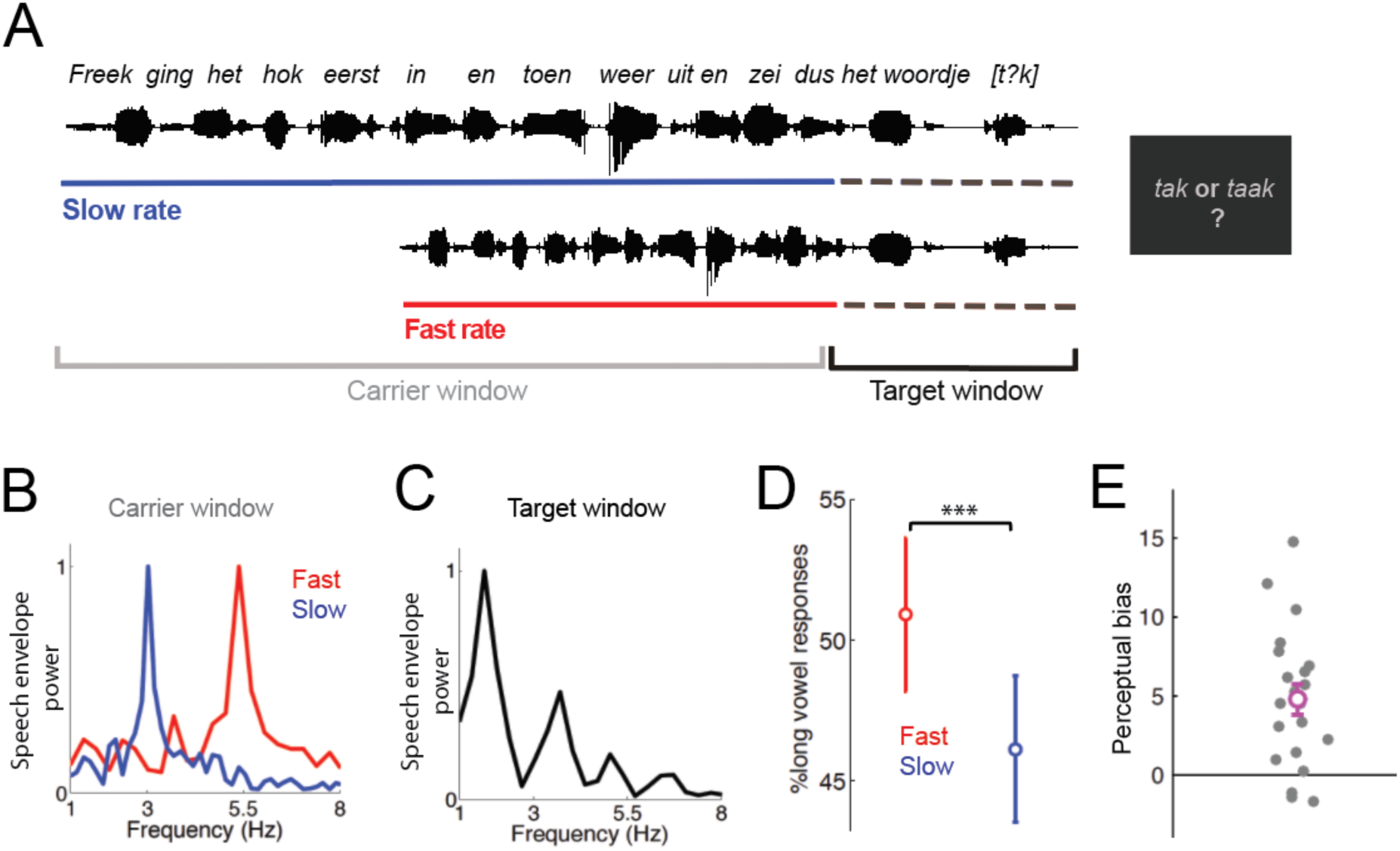
Experimental design and behavioral results. A) The participants listened to Dutch sentences with two distinct speech rates. The beginning of the sentence (carrier window) was either presented at a fast or a slow speech rate. The last three words (target window) were spoken at the same pace between conditions. Participants were asked to report their perception of the last word of the sentence (target). The words presented in the carrier window did not contain semantic information that could bias target perception and did not contain any /α/ or /a:/ vowels. B) Normalized speech envelope power spectra in the Carrier window (average across all carrier sentences). The speech envelopes showed a strong oscillatory component at 3 Hz for the Slow (blue) condition, and at 5.5 Hz for the Fast (red) speech rate condition (the two rates correspond to the syllabic presentation rate of the stimuli). C) Normalized speech envelope power spectra in the Target window (averaged across all sentence endings). 3 Hz and 5.5 Hz oscillatory components were not prominently observed in the power spectra during the Target window. D) Proportion of long vowel percepts in the Fast (red) and Slow (blue) speech rate conditions. Error bars represent s.e.m. The perception of the target word was influenced by the initial speech rate: more long vowel percepts were reported when the word was preceded by a fast speech rate. E) Perceptual Bias. We defined the perceptual bias as the difference on long vowel reports between the Fast and Slow speech rate conditions. Each grey dot corresponds to one participant. The magenta dot corresponds to the average perceptual bias across participants. Error bars represent s.e.m.

## RESULTS

### Speech perception is influenced by contextual speech rate

Target words always contained an ambiguous vowel that could either be categorized as a short /α/ or as a long /a:/ vowel. Note that the two vowels are distinguishable by both temporal (duration) and spectral characteristics (e.g. second formant frequency; F2) [22,24]. In the design, vowels were kept at a constant duration, but were presented at three distinct F2 frequencies (one ambiguous F2 value, one F2 value biasing participant reports towards short /α/ responses, one F2 value biasing participant reports towards long /a:/ responses). The F2 was varied to control for the participant’s engagement in the task, and as expected participants relied on this acoustic cue to discriminate the two vowels (main effect of F2: *F*(2,40) = 124.5, *p* < 0.001). Crucially, the preceding speech rate affected the perception of the target word (main effect of speech rate: *F*(1,20) = 24.4, *p* < 0.001). Participants were more biased to perceiving the word with a long /a:/ vowel (e.g., *taak*) after a fast speech rate, and the word with a short /α/ vowel (e.g., *tak*) after a slow speech rate (Fig. 1D). We quantified how strongly each participant was affected by the preceding speech rate in his/her behavioral report with the Perceptual Bias, which corresponds to the difference in the percentage of long /a:/ vowel reports between the Fast and Slow rate conditions (Fig. 1E). As the behavioral effect of contextual speech rate was not significantly different across the various F2s tested (interaction F2 by speech rate: F(2,40) = 0.6, p = 0.58), we pooled the data across F2 conditions for the following MEG analyses.

### Neural entrainment to the acoustics of the speech envelope during the carrier window

The MEG analysis was performed at two distinct time windows: the carrier window (sentence presentation up to the change in speech rate), and the target window (sentence endings after the change in speech rate). During the carrier window, the speech envelopes in the Slow and Fast rate conditions had a strong oscillatory component at 3 Hz and 5.5 Hz respectively. Therefore, neural entrainment was expected to peak at 3 Hz for the Slow rate condition and at 5.5 Hz for the Fast rate condition. To test this, we introduced the Entrainment Index (EI, see Materials and Methods). EI is based on the ratio of neural oscillatory power at the 3 Hz and at 5.5 Hz between the Fast and Slow conditions. EI is larger than 1 when neural entrainment to the initial speech rate is observed for both Fast and Slow conditions (i.e., stronger 3 Hz power for Slow condition and stronger 5.5 Hz power for Fast condition). Significant entrainment to speech (EI > 1) was observed during the carrier window, demonstrating that low-frequency brain activity efficiently tracked the dynamics of speech (Fig. 2A). The strongest EI was most prominently observed in auditory cortices, suggesting that primarily sensory responses accounted for the observed neural entrainment (Fig. 2A, Fig. S1A). Strong EI was observed for all participants (Fig. 2B), and effectively captured the entrainment to the actual speech rate. The 3 Hz power was relatively stronger in the Slow rate condition than in the Fast rate condition, and 5.5 Hz power was stronger in the Fast rate condition (Fig. S1B).

**Figure 2:**
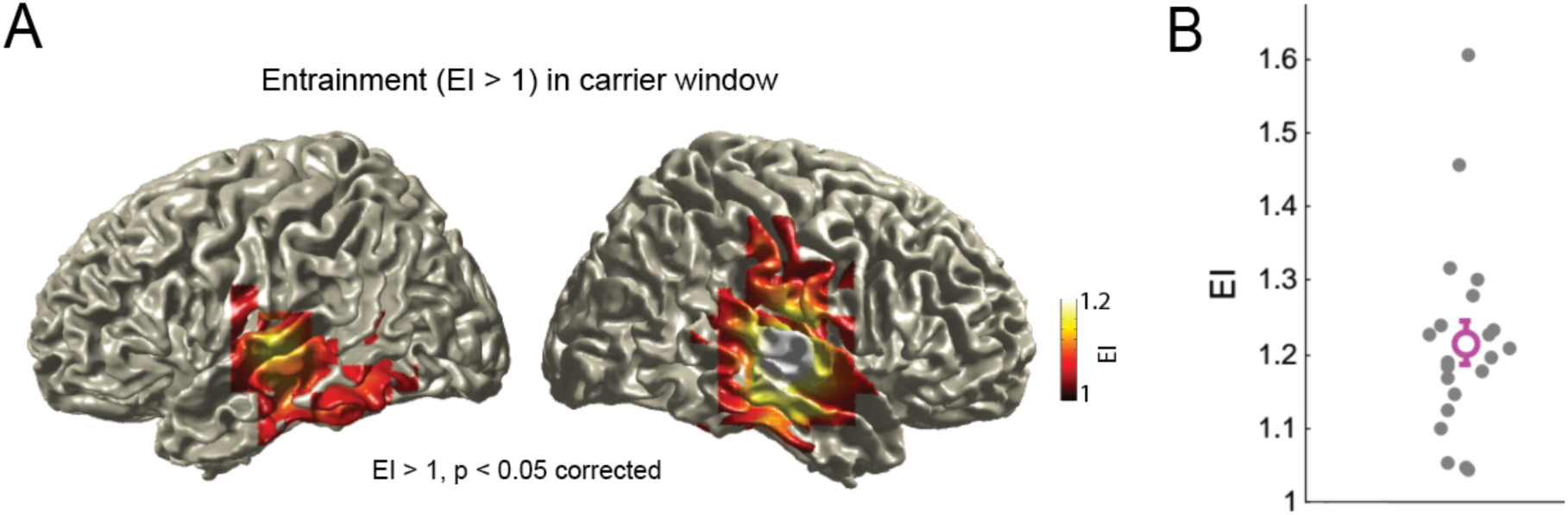
Neural entrainment to speech during carrier window. A) During the carrier window, neural oscillations in auditory areas entrain to the current speech rate (i.e. EI > 1). Top panel shows the EI values thresholded at p<0.05, controlled for multiple comparisons. B) Entrainment Index within the most strongly activated grid point (MNI coordinates: 60, −20, −10, Right Superior Temporal Cortex). Each grey dot corresponds to one participant. The magenta circle corresponds to the average EI across participants. Error bars represent s.e.m.

### Neural entrainment to past speech dynamics persists after the change in speech rate and affects comprehension

EI was also significantly larger than 1 during the target window, in which the speech acoustics were identical across Fast and Slow rate conditions (Fig. 3A-B, Fig. S2A). Larger EI (>1) reflected stronger oscillatory response that corresponded in frequency to the preceding speech rate (3 Hz power in the Slow rate condition and 5.5 Hz power in the Fast rate condition, Fig. S2B) even though the speech signals did not contain a high 3 or 5.5 Hz power (Fig. 1C), suggesting that neural entrainment to the preceding speech rhythm persisted. Sustained entrainment was most prominently observed along the right superior temporal and inferior temporal sulci, with the significant cluster extending to the right infero-frontal areas (Fig. 3A). No significant sustained entrainment was observed in the left hemisphere during the Target Window (Fig. 3A, Fig. S2 A).

**Figure 3:**
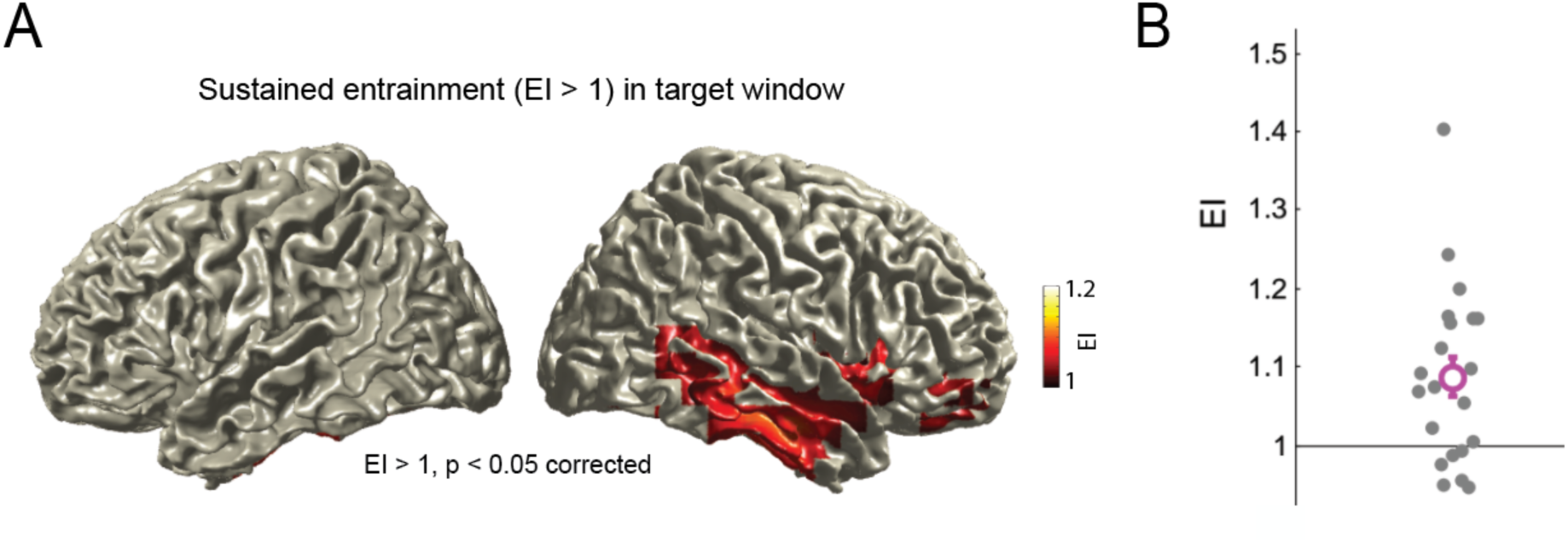
Sustained neural entrainment during target window. A) During the target window, sustained entrainment to the preceding speech rate was observed, most prominently in right middle-temporal and right infero-frontal areas. EI values are thresholded at p<0.05, controlled for multiple comparisons B) Entrainment Index within the most strongly activated grid point (MNI coordinates: 50, −40, −10, Right Middle Temporal Cortex). Each grey dot corresponds to one participant. The magenta circle corresponds to the average EI across participants. Error bars represent s.e.m.

Crucially, sustained entrainment correlated with behavioral performance, so that participants with stronger entrainment were also more strongly biased in their perceptual reports by the contextual speech rate. We correlated the EI observed in the most activated grid point of the significant cluster (in right middle temporal cortex, MNI coordinates: 50, −40, −10) to the Perceptual Bias of each participant. A significant positive correlation was observed between the two measures (Spearman’s rho: 0.54, *p* = 0.018, Fig. 4A), suggesting that participants with stronger sustained entrainment (i.e. high EI) had a stronger Perceptual Bias, i.e., were more influenced by the preceding speech rate in the perception of the target word (more likely to perceive a short /α/ after a slow speech rate, and a long /a:/ after a fast speech rate). Hence, inter-subject variability in the strength of sustained entrainment was observed and could predict how susceptible participants’ judgments on the target word were affected by contextual speech rate.

**Figure 4:**
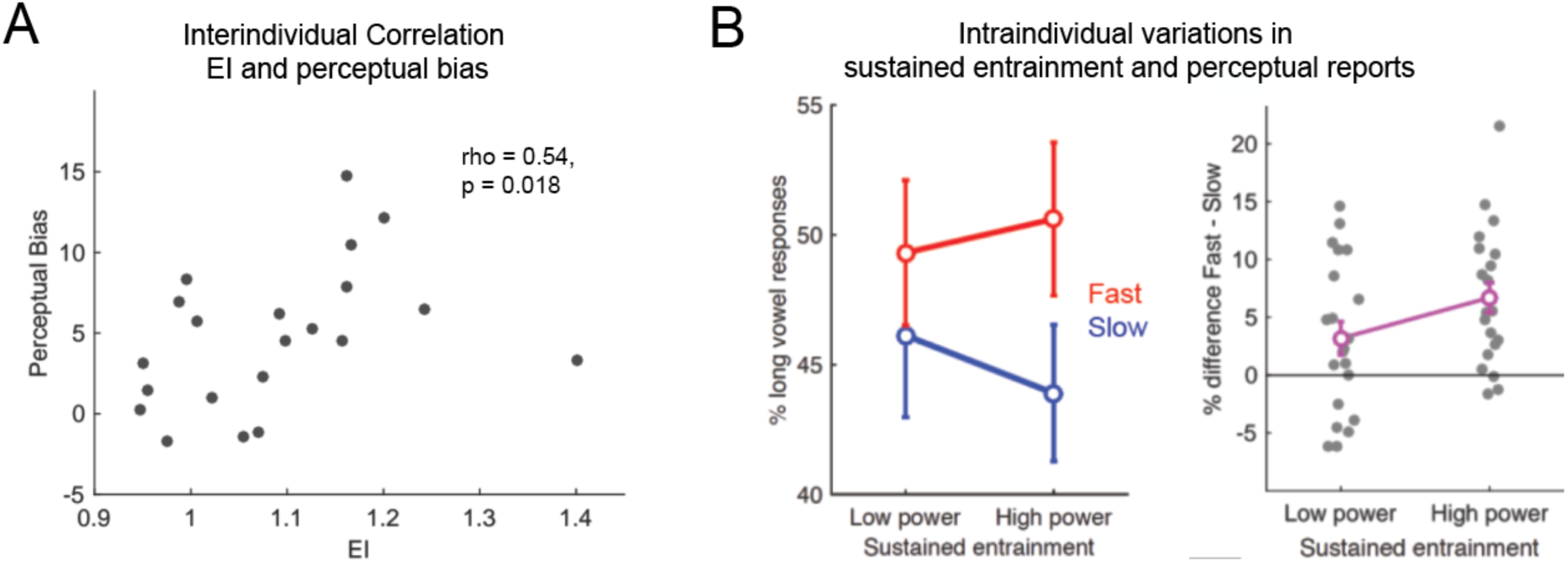
Sustained neural entrainment during target window predicts speech perception. A) Correlation between sustained entrainment (as measured by EI) and Perceptual Bias. Each dot corresponds to one participant. The stronger the sustained entrainment, the stronger the influence of preceding speech rate on target word percept. B) For each participant, the data were median split based on the strength of sustained entrainment to the preceding speech rate. For the Fast rate condition (red), the trials were split based on the observed 5.5 Hz power. For the Slow rate condition, the trials were divided based on the observed 3 Hz power. Left panel: Proportion of long vowel responses as a function of the strength of the sustained entrainment for the Fast (red) and Slow (blue) rate conditions. Error bars denote s.e.m. More long vowels percepts were observed for trials with strong sustained entrainment to the Fast speech rate; conversely more short vowel percepts were observed for trials with strong sustained entrainment to the Slow speech rate. Right panel: Perceptual Bias as a function of the strength of sustained entrainment. Each dot corresponds to one participant. The magenta circle corresponds to the average perceptual bias across participants. Error bars denote s.e.m. The stronger the sustained entrainment to the preceding speech rate, the stronger the perceptual bias.

We also asked whether sustained entrainment positively correlated with the Perceptual Bias on a trial-by-trial basis. For each participant, the individual data were split into two groups of trials based on the strength of sustained entrainment in the Target window. For the Fast rate condition, the trials were median-split based on the power of sustained 5.5 Hz oscillations. For the Slow rate condition, the trials were divided based on the observed 3 Hz power. We observed that the strength of sustained entrainment impacted the perceptual reports at the trial level. More long vowel percepts were observed for trials with strong sustained entrainment to the Fast speech rate; conversely more short vowel percepts were observed for trials with strong sustained entrainment to the Slow speech rate (Fig. 4B, left panel, interaction between Speech rate and Strength of sustained entrainment *F*(1,20) = 3.77; *p* = 0.066, marginally significant). Stronger sustained entrainment was thus associated with a stronger Perceptual Bias (Fig. 4B, right panel).

## DISCUSSION

We investigated neural oscillatory activity during listening to sentences with changing speech rates. We observed that neural oscillations entrained to the syllabic rhythm at the beginning of the sentence (carrier window). Crucially, entrainment to the preceding speech rate persisted after the speech rate had suddenly changed (i.e., in the Target window). The observed sustained entrainment biased the perception of ambiguous words in the Target window. The participants who demonstrated stronger sustained entrainment were also more influenced by the preceding speech rate in their perceptual reports. Strong sustained slow rate entrainment was associated with a bias towards short vowel word percepts, and strong sustained fast rate entrainment biased towards more long vowel percepts.

To our knowledge, the present results provide the first evidence in human recordings that neural entrainment to speech outlasts the stimulation. Sustained neural entrainment is a crucial prediction in support of the active nature of neural entrainment [1]. First, the sustained entrainment, being independent of the dynamics of the speech tokens, shows that low frequency entrainment to speech rhythms is not purely stimulus driven [25]. Second, sustained entrainment to the temporal statistics of past sensory information supports the hypothesis that neural entrainment builds temporal predictions [2,6]. Recent reports have shown that parieto-occipital alpha oscillations outlast rhythmic visual presentation [5] or brain stimulation [26]. In an electrophysiological study with monkeys, Lakatos and colleagues [4] showed that auditory entrainment in the delta band (1.6 – 1.8 Hz) outlasts the stimulus train for several cycles and argued that the reported sustained entrainment could be of crucial importance for speech processing. The current findings support this view, showing that sustained entrainment is observable in human temporal cortex and influences speech perception.

The present findings support oscillatory models of speech processing [8–10], which suggest that neural entrainment is a mechanism recruited for speech parsing. In these models, neural theta oscillations (4-8 Hz; entraining to syllabic rates) flexibly adapt to the ongoing speech rate and define the duration at which syllabic tokens are chunked within the continuous signal. Modulations in the frequency of entrained theta oscillations should then modify the discretization of the acoustics, potentially leading to distinct percepts of a spoken target word. In the present study, the observed effects of speech rate on the perceived vowel of the target word are then interpretable as a mismatch in the actual duration of incoming syllables and the predicted syllabic duration defined by the frequency of entrained oscillations [8,9,21,27,28]. A preceding fast rate would generate sustained neural entrainment of a faster rate (i.e. shorter expected syllable duration) than the monosyllabic word being parsed; this would lead to an overestimation of its duration and biasing percepts towards a word containing a long vowel. Conversely, slower sustained neural entrainment could lead to underestimation of the syllable’s duration biasing perception towards short vowel percepts. We speculate that sustained entrainment could also be at the origin of other perceptual effects of contextual speech rate: if entrainment delineates parsed tokens within continuous speech, then distinct sustained entrainment frequencies could lead to changes in the perceived word segmentation [20], and sustained entrainment could even cause the omission or certain words [19] if occurring at the phase of entrained oscillations that marks the boundary between discretized tokens.

Sustained entrainment was most prominently observed in the right middle temporal areas, while it seemed to be absent in the left temporal areas. This observation is in line with evidence that the right superior temporal sulcus is specialized in processing sound events of syllabic length (∼250 ms) [29,30], and that the tracking of the speech envelope [31–33], and of slow spectral transitions [34,35] or prosodic cues [36] are known to be stronger in right auditory than in left auditory cortices [37,38]. Our findings may further imply that the general asymmetry in speech envelope tracking during listening could originate from an asymmetry in temporal expectation mechanisms. Both left and right auditory cortices may be involved in the bottom-up tracking of acoustic features in speech, but the right temporal regions would be additionally recruited for the temporal prediction of future speech events.

The results confirm that the tracking of the temporal regularities of sounds is a neural strategy used for optimizing speech processing. Yet, the relevance of neural oscillations in building temporal predictions based on past temporal statistics may be a general property of sensory processing [39,40], in line with the idea that oscillations provide temporal metrics for perception [41,42]. Additionally, the current study was focused on the neural entrainment to the strongest rhythmic cues in the speech envelope, i.e., syllabic rhythms, operated by theta oscillations (3-8 Hz). We argue that the observed sustained entrainment would primarily influence the processing of speech acoustic features considering that theta oscillations are linked to acoustic parsing [43] and phonemic processing [44,45], while they do not seem to be involved in parsing of words in the absence of relevant acoustic cues [28]. Theta oscillations would then serve a distinct role compared to oscillations in the delta range (1-3 Hz): theta would be involved in the acoustic parsing of continuous speech into words, while delta oscillations would combine the segmented words into larger linguistic discrete structures based on procedures underlying syntactic and semantic combinatoriality [13,46–48].

In summary, the present results show neural entrainment to speech is not purely stimulus driven and is influenced by past speech rate information. Sustained neural entrainment to past speech rate is observed, and it influences how ongoing words are heard. The results thus support the hypothesis that neural oscillations actively track the dynamics of speech to generate temporal predictions that would bias the processing of ongoing speech input.

## MATERIALS AND METHODS

### Participants

33 native Dutch speakers took part in the experiment. All participants provided their informed consent in accordance with the declaration of Helsinki, and the local ethics committee (CMO region Arnhem-Nijmegen). Participants had normal hearing, no speech or language disorders, and were right handed. We excluded 10 participants who presented strong bias in their perceptual reports (<20 % or >80 % long vowel reports throughout the experiment, explicit strategies reported during debriefing); two participants were excluded due to corrupted MEG data; leaving 21 participants (14 females; mean age: 22 years old) in the analysis.

### Stimuli

A female native speaker of Dutch was recorded at a comfortable speech rate producing five different sentences, each ending with “het woordje [target]” (meaning: *the word [target]*). Recordings were divided into two temporal windows. The Carrier windows were composed of the first 12 syllables prior to “het” onset; the Target windows contained the ending “het woordje [target]”. Carrier sentences did not contain semantic information that could bias target perception and did not contain any /α/ or /a:/ vowels. Carriers were first set to the mean duration of the five carriers and then expanded (133% of original rate) and compressed (1/1.33 = 75% of original) using PSOLA [49] in Praat [50], manipulating temporal properties while leaving spectral characteristics intact (e.g., pitch, formants). The resulting Fast and Slow carriers had strong periodic components at 5.5 Hz and 3 Hz, respectively (Fig 1B). The sentence-final Target window (“het woordje [target]”) was kept at the originally recorded speech rate (i.e., not compressed/expanded). As targets, the speaker produced 14 minimal Dutch word pairs that only differed in their vowel, e.g., “zag” (/zαx/) - “zaag” (/za:x/), “tak” (/tαk/) - “taak” (/ta:k/), etc… One long vowel /a:/ was selected for spectral and temporal manipulation, since the Dutch /α/-/a:/ contrast is cued by both spectral and temporal characteristics [22,51]. Temporal manipulation involved compressing the vowel to have a duration of 140 ms using PSOLA in Praat. Spectral manipulations were based on Burg’s LPC method in Praat, with the source and filter models estimated automatically from the selected vowel. The formant values in the filter models were adjusted to result in a constant F1 value (740 Hz, ambiguous between /α/ and /a:/) and 13 F2 values (1100-1700 Hz in steps of 50 Hz). Then, the source and filter models were recombined and the new vowels were adjusted to have the same overall amplitude as the original vowel. Finally, the manipulated vowel tokens were combined with one consonantal frame for each of the 14 minimal pairs.

### Procedure

Before MEG acquisition, participants were presented with a vowel categorization staircase procedure to estimate individual perceptual boundaries between /α/ and /a:/. It involved the presentation of the target word “dat” (/dαt/) - “daad” (/da:t/) in isolation (i.e., without preceding speech) with varying F2 values (1100-1700 Hz), with participants indicating what word they heard. Based on this procedure, 3 F2 values were selected, corresponding to the individual 25%, 50%, and 75% long /a:/ categorization points. These values were used in the MEG experiment, where half of the target words contained an ambiguous vowel (F2 associated to 50% long /a:/ categorization point), a quarter of the target words with a vowel F2 associated to 25% long /a:/ responses, and a quarter target words with a vowel F2 corresponding to 75% long /a:/ responses. In the MEG experiment, stimuli included carrier sentences followed by target sequences. All participants heard the five carriers in both rate conditions in combination with all possible targets in a randomized order. Participants were asked to listen to the full sentences while fixating on a fixation cross on the screen, and to report what the target word was by button press once the response screen appeared (presented 700 ms after target offset, with the two response options presented left and right, e.g., “tak” or “taak”; position counter-balanced across participants). In total, 280 sentences were presented per Slow/ Fast speech rate condition, leading to a total of 560 trials. The experiment included 3 breaks and lasted about 75 min.

### Behavioral analysis

For every participant, behavioral responses (i.e., whether the target word contained a short or a long vowel) were registered for both Fast and Slow conditions. The perceptual bias was calculated as the difference in the proportion of long vowel (/a:/) responses between the Fast and the Slow conditions. Statistical analysis was performed with Matlab R2015a. Repeated measures ANOVA were performed using the proportion of long vowel reports and the perceptual bias as dependent variables and factors of Speech rate (Fast, Slow) and second formant frequency F2 (25%, 50%, 75% long vowel reports F2s).

### MEG analysis

MEG recordings were collected using a 275-channel axial gradiometer CTF MEG system at a sampling rate of 1.2 kHz. For source reconstruction analysis, structural magnetic resonance imaging (MRI) scans were obtained from all subjects using either a 1.5 T Siemens Magnetom Avanto system or a 3 T Siemens Skyra system. MEG data was analyzed using the Fieldtrip software [52]. MEG recordings were epoched at two distinct windows (Carrier and Target). Epochs for the Carrier window comprised the MEG recordings at the start of the sentence up to the change in speech rate (fixed 3.55 s duration for the Slow rate condition, 2.0 s for the Fast rate condition). Epochs in the Target window started after the change in speech rate and comprised the MEG recordings during the presentation of the last three words of the sentence (“Het woordje [target word]”) up to 500 ms before the response screen (the window was of 1.3s duration for both Fast and Slow conditions). Noisy channels and trials with muscle artifacts were excluded after visual inspection. An independent component analysis was performed to remove cardiac and eye movement artifacts.

The sources of the observed 3 Hz and 5.5 Hz activity were computed using beamforming analysis with the dynamic imaging of coherent sources (DICS) technique [53] to the power data. The cross-spectral density data structure was computed using Fast Fourier transform (FFT) with Hanning tapering performed at 3 Hz and at 5.5 Hz for both Carrier and Target windows. For the Carrier window, the first 500 ms of the epochs were removed to exclude the evoked response to the onset of the sentence and ensure the measure of the entrainment regime. The data was zero-padded up to 4.0 s for both conditions to match in FFT resolution. During the target window, the data was zero-padded up to 2.0 s so as to obtain more accurate amplitude estimates of the resolvable 3 Hz and 5.5 Hz signals components. The co-registration of MEG data with the individual anatomical MRI was performed via the realignment of the fiducial points (nasion, left and right pre-auricular points). Lead fields were constructed using a single shell head model based on the individual anatomical MRI. Each brain volume was divided into a grid points of 1 cm voxel resolution, and warped to a template MNI brain. For each grid point the lead field matrix was calculated. Source reconstruction was then performed using a common spatial filter obtained from beaming data from both Slow and Fast speech rate conditions. The Entrainment Index (EI) was calculated based on the source reconstructed power for each grid point according to the formula:

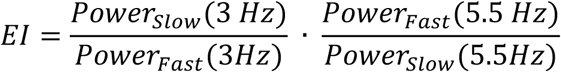

Sources with significant EI > 1 were estimated using cluster-based permutation statistics [54]. First, a “null hypothesis” source dataset was generated by setting the EI values to 1. Pairwise t-tests were then computed for each grid point between the experimental EI source data to the generated “null hypothesis” source dataset. Grid points with a p-value associated to the t-test of 5% or lower were selected as cluster candidates. The sum of the t-values within a cluster was used as the cluster-level statistic. The reference distribution for cluster-level statistics was computed by performing 1,000 permutations of the EI and the generated null hypothesis source data. Clusters were considered significant if the probability of observing a cluster test statistic of that size in the reference distribution was 0.05 or lower.

The inter-individual correlation between brain data and perceptual bias was performed within the most strongly activated grid point (grid point with highest *t*-value) located within the significant observed cluster. Single-trial power analysis was computed at this grid point to estimate the inter-trials effects of sustained entrainment on the Perceptual Bias. Single-trial time series were first computed using a Linearly constrained minimum-variance (LCMV) beamformer spatial filter. The largest of the three dipole directions of the spatial filter was kept for power analysis. The power at 3 Hz and 5.5 Hz was estimated for each trial using the same parameters as for the first analysis. The trials were sorted in two groups based on the strength of the oscillatory component corresponding to the initial speech rate (3 Hz for Slow rate condition, 5.5 Hz for Fast rate condition). The % long vowel responses were then contrasted between the two groups using a two-way repeated measure ANOVA with Speech rate (Fast, Slow) and Sustained Entrainment Strength (Low, High) as factors.

## ACKNOWLEDGEMENTS

We would like to thank Annelies van Wijngaarden for the recordings of her voice and Anne van Hoek for help with pretesting.

